# Precise, individualized foraging flights in honey bees revealed by multicopter drone-based tracking

**DOI:** 10.64898/2025.12.02.691855

**Authors:** Rachael Stentiford, Michael J.M. Harrap, Victor V. Titov, Stephan Lochner, Andrew D. Straw

## Abstract

Honey bees routinely fly hundreds or thousands of meters between their hive and established foraging locations [1–3]. To navigate these long distances, they are known to combine both landmark use [4–7] and path integration [8–11] and have been hypothesized to build a cognitive map [12–14]. Due to technical challenges inherent in tracking these small insects, obtaining three-dimensional, high-resolution measurements of individual navigational precision, and thus a detailed understanding of their strategies, has been difficult. Here, we utilize a novel multicopter drone-based tracking system [15] to measure the individual flight paths of honey bees in a structured agricultural landscape at unprecedented spatial and temporal resolution. Although bees could choose from multiple routes, we discovered that individual bees follow idiosyncratic paths with striking and repeatable precision. Flight path variability was highest over visually sparse regions and lowest near prominent proximal landmarks. Furthermore, individual strategies differed: some bees flew directly toward the hive before maneuvering around a specific tree, while others flew directly to a gap between a hedgerow and the tree. Thus, each animal varies in how it uses visual information and selects between behavioral strategies. The level of precision exhibited by their flight paths exceeds that reported in the waggle dance, implying that dance variability does not reflect a limit in the bee’s underlying spatial representation. Our results demonstrate the remarkable precision of individual bee navigation and illustrate the potential of this new drone-based tracking method to illuminate fine-scale behavioral mechanisms across a spectrum of honey bee ecology and cognition.

**Highlights:**

- Individual honey bees follow strikingly precise, idiosyncratic flight paths, revealing personalized strategies for navigating complex, obstacle-rich landscapes.
- Variability in individual bee flight paths was lower near conspicuous landmarks, but greater when further from such features, as a consequence variability in bee flights was not uniform along a route.
- Navigational precision far exceeds waggle dance variability, showing that dance imprecision is not constrained by limits in spatial representation.
- Multicopter drone-based tracking method enables new investigations of 3D flight control and visual navigation in structurally complex realistic environments.

## Results and Discussion

In an agricultural landscape, we established a honey bee hive and a feeder containing sugar water 122 meters away. In high summer, few floral resources were available and bees spontaneously learned to visit the feeder. Using retroreflective markers attached to individual bees (Figure 1A) and a drone equipped with a fast lock tracking system [15], we tracked flights between the feeder (Figure 1B) and the hive (Figure 1C). We recorded 255 full flights from 26 bees (Figure S2A). This included 92 outbound (from hive to feeder; Video S1) and 163 inbound flights (from feeder to hive; Video S2). The hive was positioned facing north under a large tree near the end of a large hedgerow. The feeder was just outside a cornfield to the south of the hive (Figure 1; corn was approximately 2 meters tall). A direct flight between the hive and feeder was blocked by a large tree. Bees were required maneuver around, or directly over, the tree. No bees were observed flying directly over the tree. Outbound bees were observed to take three different routes (Figure 1D): ‘around the tree’ where the bee flew to the west side of the tree via a gap in the hedgerow, ‘over the hedge’ where bees flew to the east of the hive and crossed over the hedgerow and around the tree via the east side; ‘wide of the hedge’, where bees travel further east traveling around the east end of the hedgerow (note that ‘route’ refers to these categorical descriptions). Of the outbound flights recorded 59 went around the tree, 28 over the hedge and 5 wide of the hedge. Inbound bees were more consistent in route choice, with 155 flights taking the ‘around the tree’ route and only 8 flights from a single bee going ‘over the hedge’. We successfully tracked some bees for only one full flight in either the outbound or inbound direction, and these were excluded from some analyses (Figure S2A).

**Figure 1.**
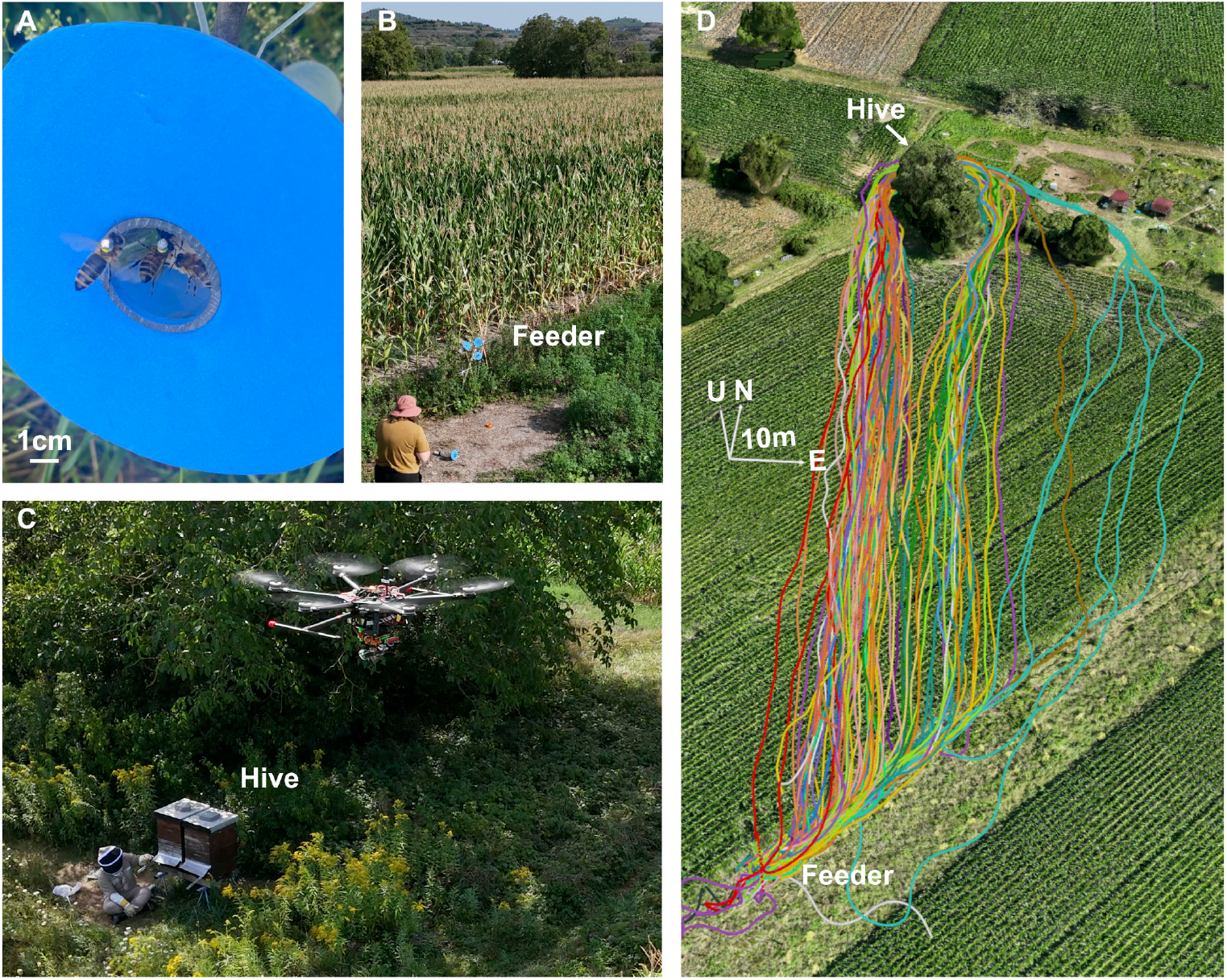
Reflector-tagged honey bees, feeder, hive and three-dimensional bee-tracking drone. (a) Reflector tags (3 millimeter diameter, 20 milligrams) are glued to the dorsal thorax of foragers captured at a feeder consisting of several artificial flowers containing sugar water. (b) The feeder is adjacent to a corn field and is shorter than the corn plants. (c) The hive is adjacent to a large tree which blocks direct southwards flight to the feeder. A multicopter drone with a fast lock-on tracking system follows bees as they emerge from the hive. (d) All outbound flights from the hive included in our analyses (n=92 flights, N=25 individuals), see Figure S1A for all inbound flights. Axes coordinates are East (E), North (N), and Up (U). Each color is one individual bee.

Out of 26 bees, 9 were observed to always take the same flight route on their outbound and inbound flights (e.g. Figure 2A), whereas 13 took different outbound and inbound routes on at least one occasion (e.g. Figure 2B, left). Five of these always used different routes between outbound and inbound legs. We observed 8 bees that took both the ‘around the tree’ and ‘over the hedge’ routes during outbound flights (e.g. Figure 2C), but there was no consistent trend in the route chosen over time (Figure S2B), suggesting the changes in outbound route choice between flights was not caused by bees learning a better route over the sampling period. We speculate that variation in outbound route choice is due to bees using a vector navigation strategy to guide this flight phase. In this view, because the ultimate goal is to reach the vector endpoint and the bee uses path integration to maintain an estimate of its own position, the specific trajectory used to bypass the tree obstacle may be irrelevant, provided the bee successfully navigates to the intended destination. Ants have also been seen to take multiple highly precise routes, apparently switching between them randomly [16].

**Figure 2.**
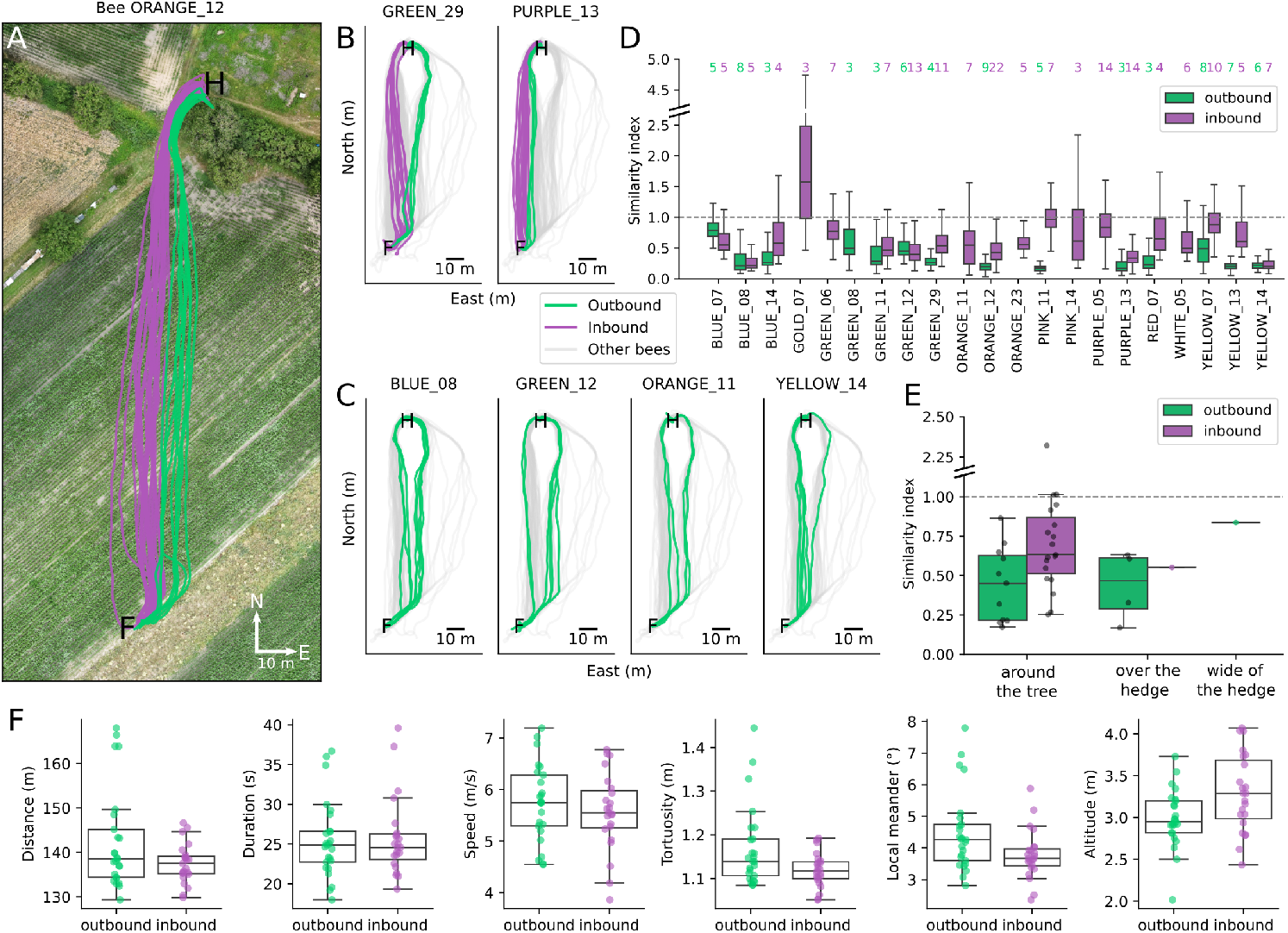
Variability of route taken. (a) Outbound (green) and inbound (purple) flights of one example bee traveling between the hive (H) and the feeder (F). (b) Further examples of outbound (green) and inbound (purple) flights of two bees that took either different routes on their outbound and inbound flights (left) or the same route on their outbound and inbound flights (right). Outbound and inbound trajectories of all other bees are also shown (grey). (c) Outbound flights (green) of four example bees that take multiple routes to the feeder. Outbound trajectories of all other bees are also shown (grey). (d) Similarity index per individual. Computation requires at least 3 trajectories per route-direction combination per bee, resulting in N = 21 bees for this similarity metric. Values below 1 indicate the trajectories of the individual are more similar to each other than to the population (high within-bee consistency). Values above 1 indicate that trajectories of the individual vary more than between bees (low within-bee consistency). Green and purple numbers indicate the total number of flights included for this bee. (e) Average similarity index by route over all bees. The most frequently taken route was ‘around the tree’ to the west. Only 1 bee made inbound flights ‘over the hedge’. Only 1 bee made outbound flights that went ‘wide around the hedge’ to the east, and no bees took this route for their inbound flights. Similarity index is significantly below 1 for outbound (one-tailed t-test, t=−7.675, df=10, p <0.001) and inbound (one-tailed t-test, t=−2.517, df=18, p = 0.011) flights around the tree and outbound flights over the hedge (One-tailed t-test, t=−5.065, df=3, p = 0.007). (f) Mean flight metrics of each bee in each travel direction (23 inbound and 25 outbound flights), calculated accross all trajectories (92 outbound, 163 inbound). No difference between outbound and inbound flights for flight metrics: distance (t-test, t=−2.00, p = 0.052), duration (t-test, t=0.16, p = 0.876), speed (t-test, t=−0.83, p = 0.413). Tortuocity of outbound flights was higher than inbound flights (t-test, t=−2.41, p = 0.020). Local meander of outbound flights was higher than inbound flights (t-test, t=−2.28, p = 0.028). Mean altitude of outbound flights is significantly lower than inbound flights (t-test, t=2.56, p = 0.014). Degrees of freedom for all tests associated with flight metrics are 46.

That the same bees can take different routes across multiple foraging flights between the hive and the feeder shows there can be large variability in the flights taken by an individual bee. However, when we focus on the trajectories within each route separately, we find strikingly high consistency in individual bee trajectories (Figure 2D,E). This result at the landscape scale is consistent with results showing that bumble bees learn idiosyncratic routes in cluttered artificial environments at the meter scale [17] or ants walking in natural habitats [16,18–20]. Low within-route variability on the outbound route may be achieved if the bees perform a consistent maneuver around the tree followed by consistent vector-guided flight. Inbound flights, while still highly consistent, had higher variability than outbound flights. This may be due to the limited visual structure available at the start of the inbound flights. In distance, duration and speed flight metrics, the outbound and inbound flights did not differ (Figure 2F). Bees flew approximately 138 meters in around 25 seconds (about 5.5–6 m/s) regardless of direction. Measures of flight straightness (tortuosity and local meander) showed outbound flights were less straight than inbound flights. The average altitude of outbound flights was also lower than inbound flights, which may be accounted for by bees taking a route where they were able to fly lower in some sections whilst following the edge of the corn (Figure 1D).

We largely observed high within-bee consistency. The exception, bee GOLD_07, had inconsistent inbound flights over the cornfield but, like other bees, converged tightly at the west side of the tree before the hive (Figure S2A). GOLD_07 also performed the only outbound flight recorded that overshot the feeder (Figure S1B). Intriguingly, in this failed flight, GOLD_07 took a different route (‘wide of the hedge’) compared to the previously recorded outbound flight (‘over the hedge’). Perhaps suggesting GOLD_07 failed to find the feeder due to an error in estimating position. We note that this flight displayed prominent oscillations after the bee failed to find the feeder, consistent with exploratory search behavior. Oscillatory paths are well documented in walking insects [21–23] (see [24] for review). Similar ocillatory looping search patterns have been seen in flying bees in previous lab [25,26] and field settings [27,28], including honey bees tracked with harmonic radar [29]. Our observations of GOLD_07 suggest that oscillations may also be important for exploratory or search behavior in bees and warrant further investigation. Computational and robotic models indicate that oscillations can be useful beyond exploration, such as in visual route following by improving view matching, familiarity-based guidance, and optic flow for altitude control [30–36]. However, the oscillations in this one outbound flight of GOLD_07 are a notable exception compared to the majority of flights in which oscillations are much less prominent.

We next examined variability along each route to understand where flight trajectories were most precise. For each bee, we computed the mean trajectory for each route and direction combination the bee took. We then defined planes perpendicular to this mean trajectory at 1 meter intervals and found the points where each trajectory intersected each plane. We then measured the spread of intersection points within each plane as the distance from the within-combination median intersection point. Variability was not constant along trajectories following the same route and variability across the route differed depending on direction of travel (Figure 3A, 3B). For ‘around the tree’ outbound flights, variability grew steadily over the flight and peaked at plane 109 near the edge of the cornfield. Near the start of inbound flights for this route, the variability increased sharply (local maximum at plane 109) and remained high (absolute maximum at plane 64) and then rapidly decreased near the end of the flight. The area of highest precision coincided with proximity to the tree in both outbound flights (plane 2) and inbound flights (plane 6), while areas of higher variability occurred over the cornfield in both flight directions. At planes with low variability, intersection points are clustered tightly (Figure 3C, top). Across all bees, intersection points at plane 6 averaged within 0.5 meter of the median intersection point (Figure 3D). This suggests the presence of conspicuous proximal landmarks such as the tree may contribute to regions of extreme flight precision. Likewise, precision in navigation may be lower when the proximal environment is more uniform and the only available landmarks more distant, such as over a highly repetitive cornfield. Proximal and distal landmarks have been shown to be utilized by bees in navigational tasks [37–43]. Information from proximal landmarks are often prioritized by honey bees over distal landmarks, and proximal landmarks may be more informative of position [44–46].

**Figure 3.**
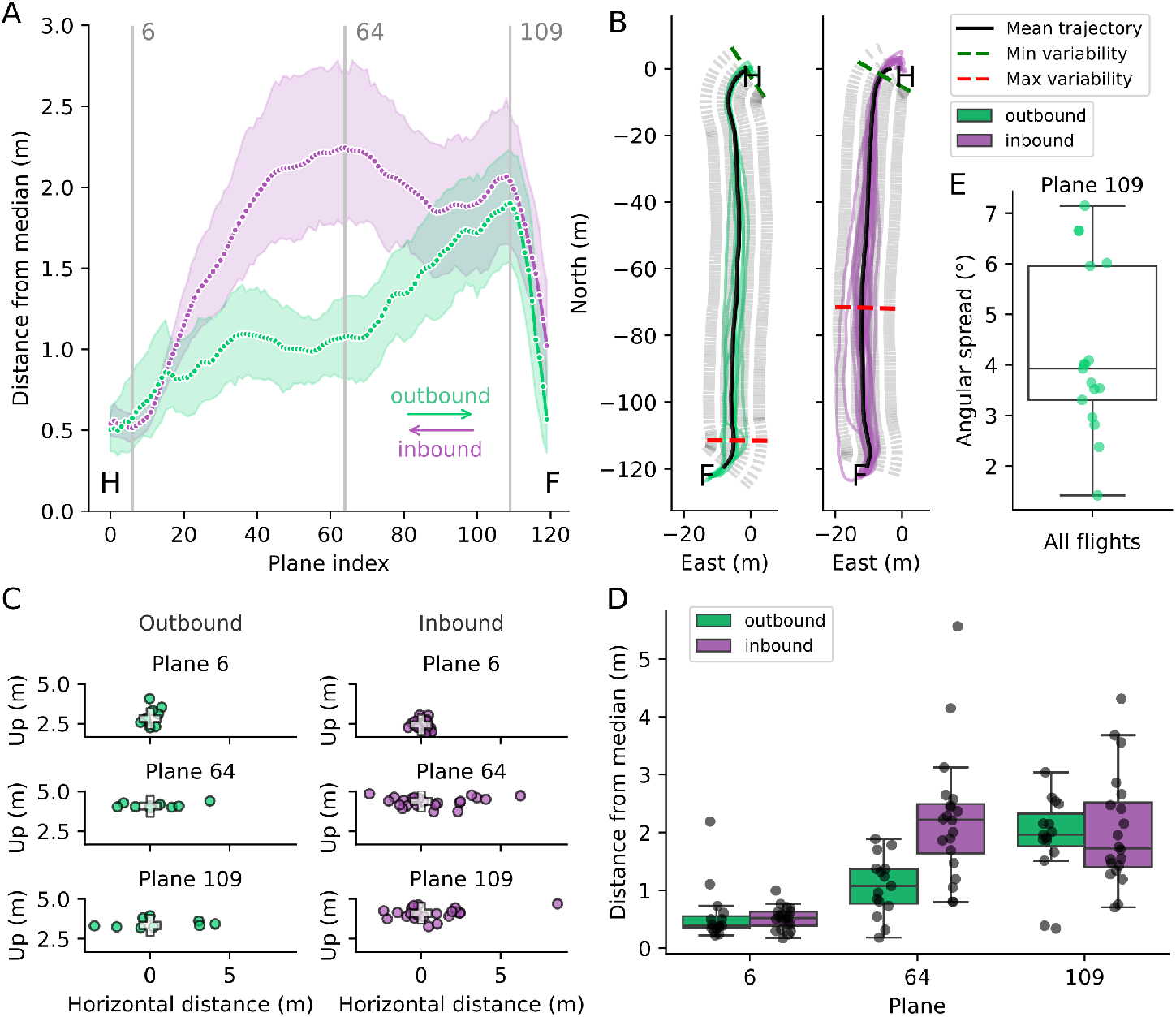
Variability along the flight path is related to landscape structure. (a) Variability of the outbound (green) and inbound (purple) flights measured at 1m increment planes along the ‘around the tree’ route (±95% confidence intervals; see Methods). Plane 0 is at the hive (H) and plane 120 just before the feeder (F). Lowest variability is at plane 2 for outbound and 6 for inbound flights. Highest variability is found at plane 109 for outbound and 64 inbound flights. (b) Outbound (left) and inbound (right) trajectories of one example bee (ORANGE_12), showing the plane with minimum variability (green; plane 0 outbound; plane 2 inbound) and maximum variability (red; plane 111 outbound; plane 71 inbound). The average trajectory for which the tangential planes were found is shown (black) along with all 120 planes (grey dashed lines) specific to each averaged route. (c) Intersection points of all trajectories of one example bee (ORANGE_12) with 3 planes: plane 6 (top), plane 64 (middle), plane 109 (bottom), for outbound (left) and inbound (right) flights. White crosses indicate the median intersection points for the ‘around the tree’ route. (d) Variability of outbound and inbound flights of the ‘around the tree’ route at 3 planes (6, 64, 109). Black points indicate variability for each bee. (e) Angular spread, measured from the hive, of intersection points at plane 109 for all flights from all three outbound route categories, each individual point is the angular spread measured per bee, showing a mean angular spread of 3.30° (±1.79° standard deviation) across the population.

The gradual accumulation of variability observed over the outbound flights (Figure 3A) is consistent with bees using a vector-based navigation strategy for this phase of the outbound flight [47]. We speculate that this strategy may be used over the cornfield, from where the feeder is invisible and the bee has no conspicuous proximal landmarks but only repetitive local visual cues. Honey and bumble bees have been shown to follow linear ground features such as paths or roads, especially when prominent skyline cues are absent [28,48–50]. Immediately upon passing the cornfield boundary, outbound trajectories often turned towards the feeder following the field boundary (Figure 1D, and 2A), contributing to decreasing variability in the final meters prior to reaching the feeder. In contrast to bumble bees that eventually abandon such features for more direct paths [28], our bees consistently followed field boundaries over repeated flights. Pheromones released from the Nasonov gland of previous foragers may have contributed to the final phase of feeder localization [51]. When flying back to the hive, the need to gain altitude above the corn before making progress toward the hive may explain the sharp rise in variability at the start of inbound flights (Figure 3A). The profile of variability in inbound flights may reflect bees utilizing a more landmark-based navigation strategy, heading towards a conspicuous landmark associated with the hive (the tree and hedgerow), as opposed to a particular vector, before correcting position to follow a precisely learned path once landmarks become more proximal (as in [38]). Individual differences may arise from differences in neural wiring, for example in the (visual) projection neuron input to mushroom body Kenyon cells [52] or earlier in the visual system [53]. Overall, these findings suggest that landscape structure, specifically the availability and location of proximal landmarks, influence the navigation strategy employed by bees.

Navigational precision of honey bees has also been investigated in the waggle dance, where it has long been recognized that precision could be under genetic, and thus evolutionary, control [54]. The tuned-error hypothesis suggests that variability in dances is likely adaptive to tune precision of recruit localization to resource distribution [55,56]. More recent work, however, argued that waggle dance precision is limited by internal constraints of the dancer [57–60]. We found honey bees could regularly replicate the same paths to high precision over multiple flights. Even at the plane of highest variability, intersection points were on average within 2 meters of the median, equivalent to an angular spread of approximately 3.30° (±1.79° standard deviation) relative to the hive (Figure 3E). This is much lower than the angular divergence observed in waggle dances for resources at similar distances (about 30° at 100 meters [55]). The low within-bee variability and small angular spread measured in the flight trajectories suggest that navigational capacity does not constrain waggle dance precision. Consequently, our data indicate that dance variability arises from either an adaptive benefit or constraints specifically in dancing.

Observing lowest variability to be west of the hive, we asked whether inbound flights travel directly towards the hive and then divert around the tree, or travel directly to the gap in the hedgerow. We found the direction of travel from the trajectory points and then compared the angle between this heading and the heading towards either the hive or the gap for inbound flights, and towards the feeder for outbound flights. During inbound flights, the proportion of the flight aligned to the hive was lower than the proportion of the flight aligned to the gap across a range of angular thresholds (Figure 4A). When flights were divided into 20 segments, alignment to the hive was higher early in the flight and decreased toward the end (Figure 4B,C). When looking at individual inbound trajectories, it seems that while each individual bee repeats a rather consistent path, the navigational strategy employed may vary between individuals (Figure 4D). Bees that fly initially towards the hive (such as BLUE_07) may initially perform vector navigation and wind down their path integration vector before making an avoidance maneuver around the tree (Figure 4D left), while others (such as ORANGE_12 and YELLOW_14) fly directly to the gap (Figure 4D middle and right), suggesting a visually guided path based on learned landmarks. Those using the direct-to-hive strategy may rely on global cues such as polarization patterns [61,62], whereas bees flying to the gap may have adopted a learned shortcut. For outbound flights, most bees undershot the feeder and then followed the field edge. This resulted in a much lower proportion of the flight aligned to the goal when compared with inbound flights. As we only tracked experienced foragers, we cannot address if the direct-to-hive strategy of some inbound flights or undershooting of outbound flights are learned strategies.

**Figure 4.**
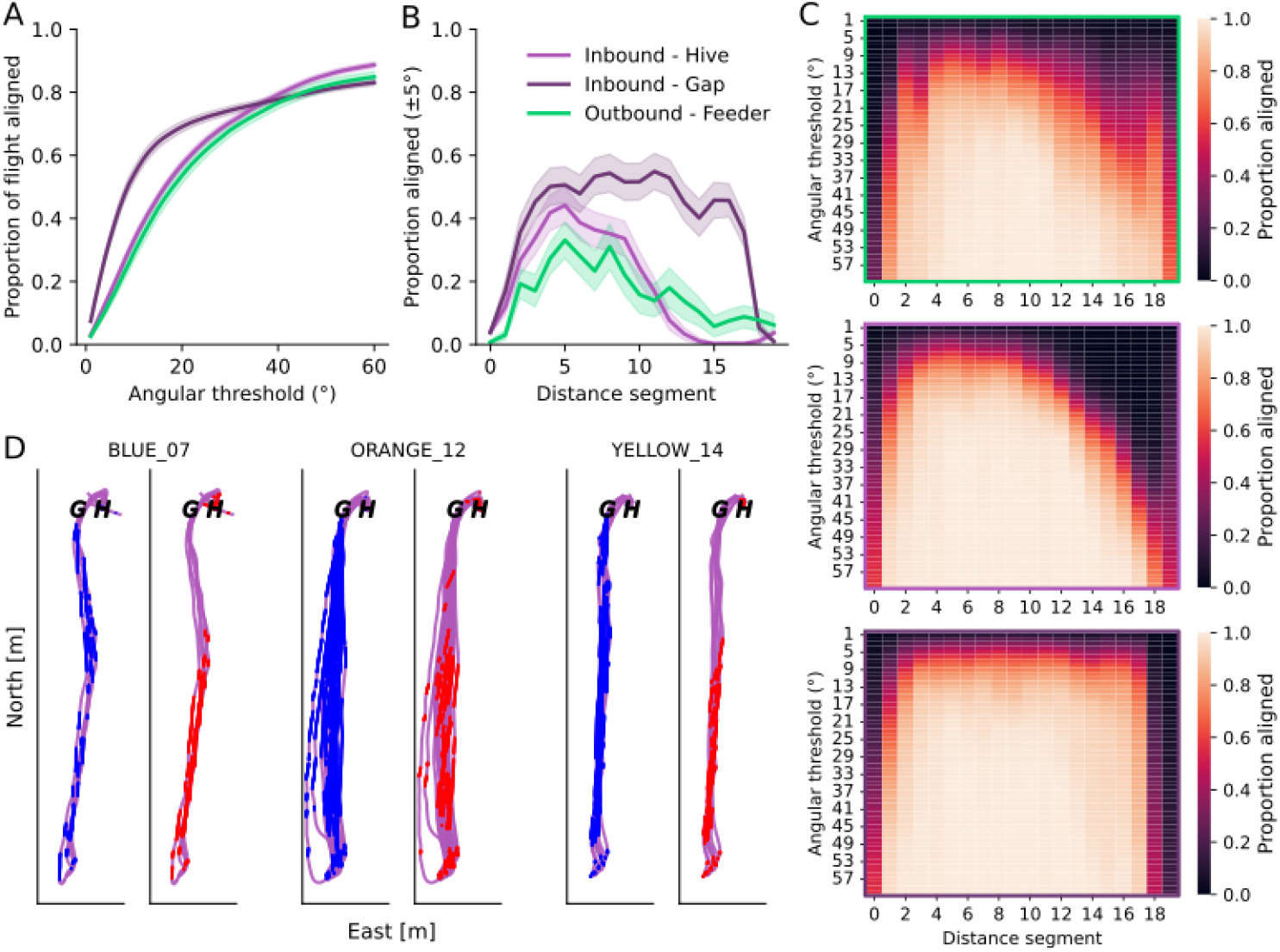
Alignment of flight direction over entire flight path. (a) Proportion of the flight aligned to within an angular threshold of a target position (±95% confidence intervals). For outbound flights this is the position of the feeder. For the inbound flights this target is either the hive or a point 8 meters west of the hive which is approximately the position of the gap between the tree and the adjacent field’s hedgerow which many bees move through when taking the route around the tree. (b) Proportion of flight aligned to within 5° of the target for each of the three targets (±95% confidence intervals). (c) Heat maps showing the proportion of flight aligned to each of the three targets: Outbound - Feeder (top), Inbound - Hive (middle), and Inbound - Gap (bottom), at different angular thresholds when the trajectories are divided into 20 segments. (d) Inbound flights for three example bees. Trajectory points aligned within 5° to the hive (H) are shown in red and trajectory points aligned within 5° to the gap (G) are shown in blue.

Here we used a recently developed multicopter drone-based bee tracking system [15] to record high resolution 3D trajectories of bees at landscape scale within a structured agricultural environment. This drone-based tracking system overcomes limitations of harmonic radar-based bee tracking [63] in that it isn’t limited by the necessity to mechanically sweep a narrow radar beam in a circle, causing low temporal resolution and 2D-only output, and while both systems are blocked by vegetation and other structures, the drone can maneuver around occlusions. These limitations of radar tracking have necessitated studies conducted in flat, open environments largely lacking vertical structure [13,64– 67]. Studies in less open environments typically cannot use radar tracking and often rely on spatially sparse information obtained by chasing marked animals [68], monitoring ‘vanishing direction’ [12,69], or mark-recapture methods [48,70]. In this work, we tracked bees in a landscape which would have precluded radar-based tracking. In doing so, we demonstrate how landscape structure influences behavior of bees, necessitating maneuvers around such obstacles when direct flight is not possible and informing the navigation strategies utilized. The drone-based tracking system is also capable of tracking bees over larger scales (see Figure S2C). We can now ask questions of how bees overcome navigational challenges in rich and complex environments, in three dimensions, where direct vector flight is not possible and tracking must be conducted around and over large features that would serve as obstacles to radar.

Our findings demonstrate that individual bees follow highly precise, idiosyncratic flight paths through complex, three-dimensional environments. Variability was lowest near major landmarks and highest mid-flight (over the cornfield), consistent with use of visual landmarks to anchor navigation and greater uncertainty in visually featureless areas. Navigational precision of experienced foragers far exceeds published waggle dance divergence at comparable distances, indicating that dance imprecision is unlikely to reflect limits in bees representation of space. Finally, inbound flights revealed two potential navigation strategies: Some bees fly directly toward the hive, then perform an avoidance maneuver around the tree. Others fly directly to a gap in the hedge, suggesting a visual landmark– based shortcut. This suggests individual differences in navigation strategy. Multicopter drone-based tracking enables new investigation of 3D flight control, obstacle negotiation, and visual navigation in realistic settings.

## Resource availability

### Lead contact

Requests for further information and resources should be directed to and will be fulfilled by the lead contact, Andrew D. Straw.

### Materials availability

This study did not generate new unique reagents.

### Data and code availability

- Trajectory estimates and code used to make all figures, including instructions required to reanalyze the data, will be made publicly available at Zenodo upon publication.
- Any additional information required to reanalyze the data reported in this paper is available from the lead contact upon request.

## Acknowledgements

This work was funded by the VolkswagenFoundation Momentum Program (98 692 to A.D.S.) and an Deutsche Forschungsgemeinschaft grant (543356743 to A.D.S.).

We thank M. Siegel and the mechanical workshop of the Institute of Biology I. The Pietsch and Suntz families kindly provided the field site. M. Tritschler, C. Hitzfeld, N. Brehm, and T. Albonetti provided bee keeping services. Data for environmental reconstruction was provided by T. Kattenborn and M. Quinten.

## Author contributions

Conceptualization, R.S., M.J.M.H., and A.D.S; Data curation, R.S. and S.L; Formal analysis, R.S. and A.D.S; Funding acquisition, A.D.S.; Investigation, R.S., M.J.M.H., V.V.T., and A.D.S.; Methodology, R.S., M.J.M.H., V.V.T., and A.D.S.; Software, R.S., S.L, V.V.T., and A.D.S.; Writing – original draft, R.S. and A.D.S.; Writing – review & editing, R.S., M.J.M.H., and A.D.S.

## STAR★Methods

### KEY RESOURCES TABLE

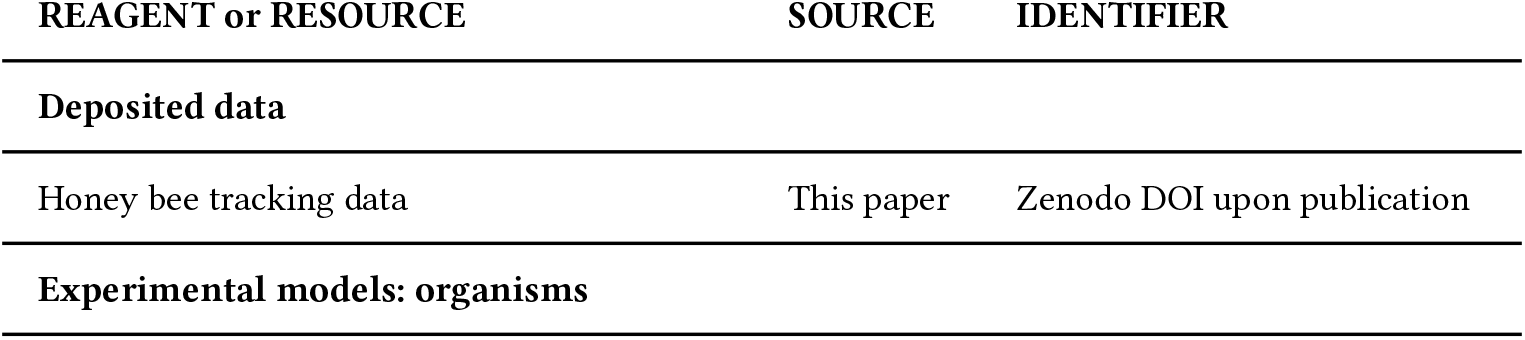
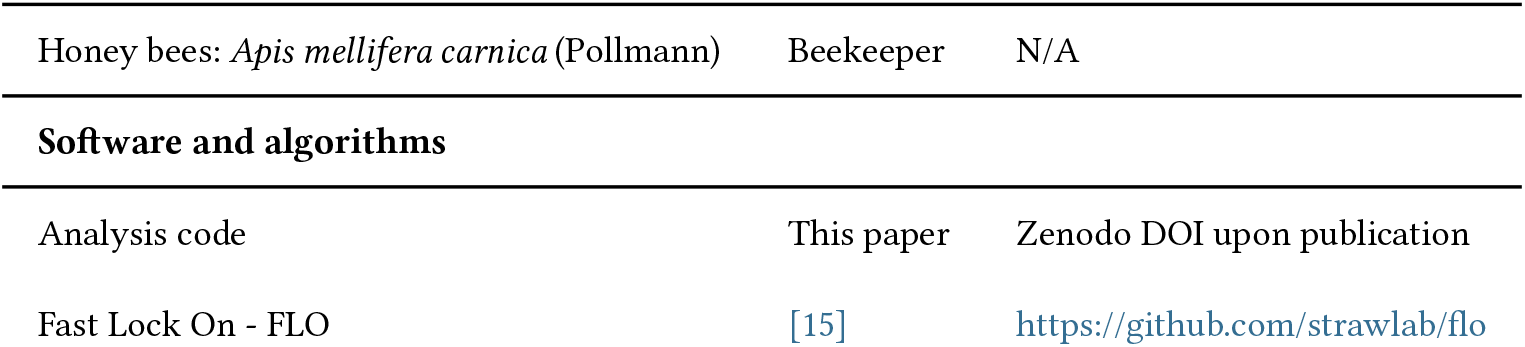

| 231 | REAGENT or RESOURCE | SOURCE | IDENTIFIER |
| --- | --- | --- | --- |
| 232 | **Deposited data** |  |  |
| 233 | Honey bee tracking data | This paper | Zenodo DOI upon publication |
| 234 | **Experimental models: organisms** |  |  |

| REAGENT or RESOURCE | SOURCE | IDENTIFIER |
| --- | --- | --- |
| Honey bees: *Apis mellifera carnica* (Pollmann) | Beekeeper | N/A |
| Software and algorithms |  |  |
| Analysis code | This paper | Zenodo DOI upon publication |
| Fast Lock On - FLO | [15] | https://github.com/strawlab/flo |

## EXPERIMENTAL MODEL AND SUBJECT DETAILS

### Honey bees and hive

Two honey bee (*Apis mellifera* L., subspecies *carnica*, Pollmann) hives were positioned facing north behind a field boundary formed of trees and hedges (Figure 1C) in the area of Ihringen am Kaiserstuhl, Germany (48.036075°N, 7.630252°E). These hives were maintained by standard beekeeping practices, and left to forage on local flora. The hive entrances were fitted with a bi-directional funnel system to separate the flow of bees into those entering and exiting the hive. On the entrance side, a large opening on the outside funneled bees towards a 2 centimeter diameter tube extending 30 centimeter into the hive, on the exit side, a large opening inside the hive funneled bees towards a 3 centimeter diameter tube (Grünke Acryl, Schwelm, Germany) extending outside of the hive by 30 centimeter. The external tube had slots cut at 5 centimeter intervals to allow a transparent plastic gates to be added to restrict the flow of bees. It was not possible to assess the exact age of bees. No permission was required for the experiments, but the experiments were conducted according to ASAB/ABS guidelines. Experiments were performed from 23 July to 14 August 2025 with bees exclusively from the eastern of the two hives.

## METHOD DETAILS

### Feeder

The feeder was composed of five artificial flowers arranged near each other (within one meter). Each artificial flower contained a sucrose water solution. The feeder was positioned on a wildflower strip, largely devoid of flowers during the experiments, near the boundary of a corn field. The corn was approximately 2 meters tall at time of the experiments. The feeder was 122 meters south of the bee hives. The entrance to each artificial flower was made from an 11 centimeter diameter blue disk (made from rubber foam, efco creative GmbH, Rohrbach, Germany) with the entrance tube (3 centimeter diameter, see above) at the centre (Figure 1A). A wick made of cotton wool was positioned inside the artificial flower with one end accessible from outside and the other inside the sucrose reservoir. Artificial flowers were positioned at variable heights between approx. 50-80 centimeter on 2 centimeter diameter bamboo stakes. Artificial flowers were refilled on work days and were typically emptied by the bees by evening. Each morning prior to tracking, we attempted to identify bees which were visiting the empty artificial flowers with the hypothesis that they were experienced, regular visitors and would likely begin frequent foraging trips even if the sugar concentration was low enough to prevent recruitment of many new bees. This was consistent with our experience and we mostly used a sucrose concentration of 0.25 M. If too few bees were visiting the feeders, we increased recruitment by using 0.5 M solution.

### Reflector attachment

Honey bee foragers were caught as they exited the artificial flower and tagged with retroreflective markers (Figure 1A, see also [15]). These markers were made of polystyrene balls (TEDi GmbH & Co. KG, Dortmund 320 Germany) wrapped in 20 x 3 millimeter strips of flexible retroreflective tape (3M Scotchlite 8850 Reflective Material Silver marking film, 3M Company, Maplewood, Minnesota, USA) and adhered with superglue (Loctite original, Henkel Corporation, Westlake Ohio, USA) to queen marking plates (Opalith, Lyson Beekeeping, Klecza Dolna, Poland) painted with an identifying color. (The retroreflective tape has been discontinued but Scotchlite 7610 is a similar product.) Markers averaged 3 millimeter diameter and 20 milligrams (of which 3 milligrams is the queen marking plate) and were prepared in advance of bee capture. To attach a marker, the bee was first immobilized in a queen marking tube (Josef Muhr Beekeeping, Prakenbach, Germany) and the hair on their upper thorax removed. The marker was then attached to the upper thorax with a dot of superglue applied with a pipette tip. Bees were then released back into the artificial flower. After marking, bees typically resumed foraging within a few minutes. Only bees which we observed on at least one new foraging bout within the hour were tracked. This was the majority, but not all, of bees marked.

We repeately observed foragers with markers returning to the feeder or emerging from the hive for multiple days after marking, suggesting the markers had limited impacts. We also observed some reflectors fall off inside the artificial flower, so it is likely some reflectors were also lost within the hive. Where possible, we removed reflectors from bees we were no longer interested in tracking. Despite marker loss, we prevented tracking the same bee twice because we avoided marking any bee with a previously shaved thorax.

### Fast lock tracking drone

We used a multicopter drone (PM X6 Carbon Hexacopter, Premium-Modellbau, Meersburg, Germany) equipped with a fast lock on tracking system [15]. This system functions by tracking light reflected from the bee-carried marker. Onboard image analysis steers a gimbal on the drone to maintain the reflector in the field of view of two cameras. A newly developed autochase function allowed the drone to chase the bee with only safety-relevant supervision from the pilot and manual control before and after tracking lock.

### Tracking

Experimenters positioned at the hive and the feeder watched for bees with reflectors and restricted their exit using the plastic gates at the hive, or a cap on the end of the entrance tube at the artificial flowers. Care was taken to allow only one marked bee into each artificial flower, or the hive exit tube, at a time. In the case where more than one bee was present either at the hive entrance or in another artificial flower, their reflector was shielded from the sight of the drone with an additional cap on the flower entrance, a cloth or gloved hand. First a reflector was shown to the drone to establish an initial lock-on direction, this reflector was then covered, and the bee released. As the bee exited the hive entrance or artificial flower the reflector became visible and tracked automatically by the drone using autochase (Video S3). In some instances the reflectors became dirty and their reflective properties sub-optimal for tracking resulting in the loss of the bee during tracking. In these cases the bees were recaught when next seen in a bee marking tube and washed with a damp cotton bud, or the reflector was replaced with a new one. In some cases the bee was not tracked for the full flight between the feeder and the hive, flights tracked for less than 90% of the full distance were excluded from further analysis as were flights that exceeded 200m in length prior to arriving at the feeder. We also tracked some bees from the hive unexpectedly towards an unknown target to the west of the hive, in one case for approximately 600 meters before the bee crossed a trainline and was lost (Figure S2C).

### Environmental reconstruction

A DJI L2 camera on a DJI Matrice 300 drone was used to collect images which were processed with WebODM 2.9.4.

## QUANTIFICATION AND STATISTICAL ANALYSIS

### Trajectory reconstruction

Combining stereopsis-measured distance, gimbal angles, and drone pose estimated by the flight controller (PX4 Firmware v1.15.2), we could reconstruct the bee position. RTK GNSS correction signals were provided by a base station (Holybro H-RTK F9P Basis GPS RTK Modul, Premium-Modellbau, Meersburg, Germany). See [15] for full details and software. Briefly, the centroid of the brightest points in the images captured by two IR cameras is detected and used to find the distance of the bee from the cameras. The reflector position is transformed into a world coordinate system by following the kinematic chain. First using the camera calibration matrix and the distance of the centroid, position is converted from image to camera coordinates. Then these points are transformed into the drone coordinate system using the translations and rotations of the gimbal relative to the drone. These new coordinates are further transformed to the world coordinate system using the drone’s pose. Local east, north, up (ENU) world coordinates can be further transformed into latitude and longitude using the known GNSS offset and reference position. Reconstruction of a stationary marker at the start of each data collection day resulted in average root mean squared error of 31 centimeters (± 22 centimeter standard deviation) of all the points in each trajectory from that trajectory’s mean point.

### Similarity index

Hausdorff distance [71,72] is the maximum of all the distances from a point in one trajectory to the closest point in a second trajectory. Similarity index [17] is the ratio between the standard deviation of within-bee Hausdorff distances and across-bee Hausdorff distances for an equal number of randomly selected trajectories from the population (bootstrapped 1000 times). Trajectories were compared within a route and direction, requiring a minimum of 3 trajectories per group for inclusion, and plotted per bee averaging across routes, and per route with the average for each bee plotted. A value less than one for the similarity index means that trajectories from one individual are more similar to themselves than to those of group as a whole (high within-bee consistency). A value greater than one for the similarity index indicates low within-bee consistency.

### Flight metrics

For each trajectory, flight distance, duration, speed and altitude were extracted from trajectory reconstructions. Tortuocity was calculated as the ratio between actual flight distance and the straight-line distance between the hive. Local meander was calculated as the mean absolute difference between the headings of adjacent 1 meter flight segments, as used to measure the wiggliness of ant trajectories [73]. Each of these metrics were calculated for all recorded trajectories. To account for the differing number of recorded trajectories accross bees, for plotting and analyses, the mean of each metric for each bee in each travel direction was calculated.

### Trajectory averaging

For each bee, the average trajectory for its outbound and inbound flights for each route is calculated (at least 3 trajectories required for averaging). Averaging is done by first defining a reference line between the hive and the feeder. Points in each trajectory are then projected onto this line by finding the closest point on the reference line. These points are then grouped into 1 meter bins and averaged. This method minimises the influence of points which extend more north above the hive or south below the feeder. Our precision estimates therefore primarily reflect segments where the tracking lock was sustained.

### Variability

To measure the variability of the flights at incremental distances along the route, we first find for each 1 meter increment the perpendicular vertical plane to the averaged trajectory. Planes were numbered sequentially from the hive to the feeder such that bees travelled from plane 0 to 120 in outbound flights and 120 to 0 in inbound flights (Figure 3). We then find the intersection points of each trajectory with these planes. For each route and direction (outbound or inbound), the distances of each point to the median intersection point are found and averaged as a measure of variability.

### Angular spread

Angular spread is measured by finding the angle between the two furthest separated intersection points, measured from the hive.

---

**Figure S1.**
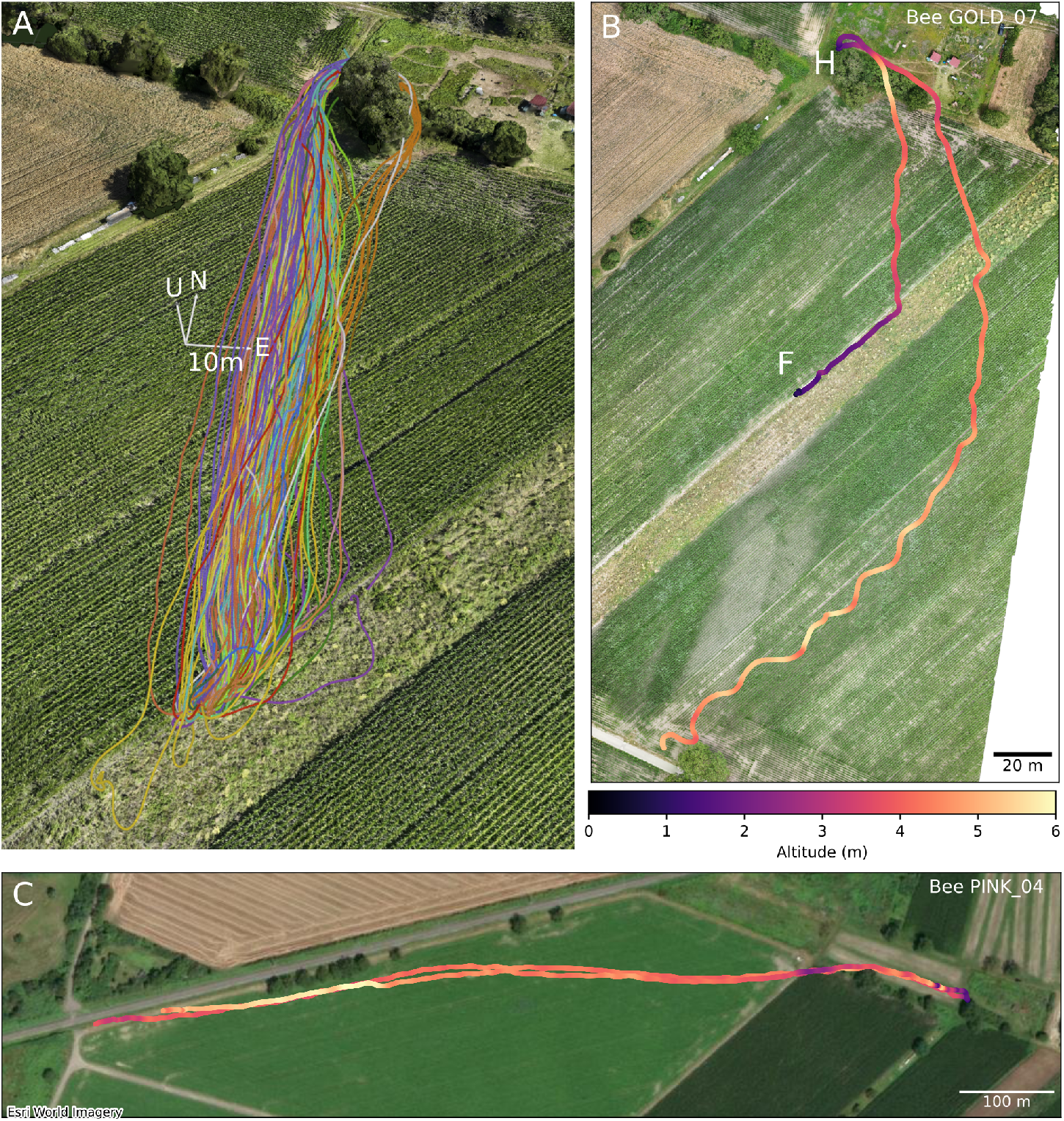
Further 3D flight paths recorded at the study site. (a) All inbound flights from the feeder included in our analyses (n=163 flights, N=26 individuals) plotted in 3D reconstruction of the study site. Axes coordinates are East (E), North (N), and Up (U). Each color is one individual bee. (b) The two recorded outbound flights of bee GOLD_07. Note how the bee successfully navigates from the hive to the feeder on one flight, but fails to reach the feeder on a subsequent attempt. (c) Two example long flights made by a bee away from the hive but not towards the feeder. The bee was tracked on both occasions until the bee reached the train tracks where tracking was no longer possible. In (b) and (c) shading of flight path indicates flight altitude with colorbar applying to both panels.

**Figure S2.**
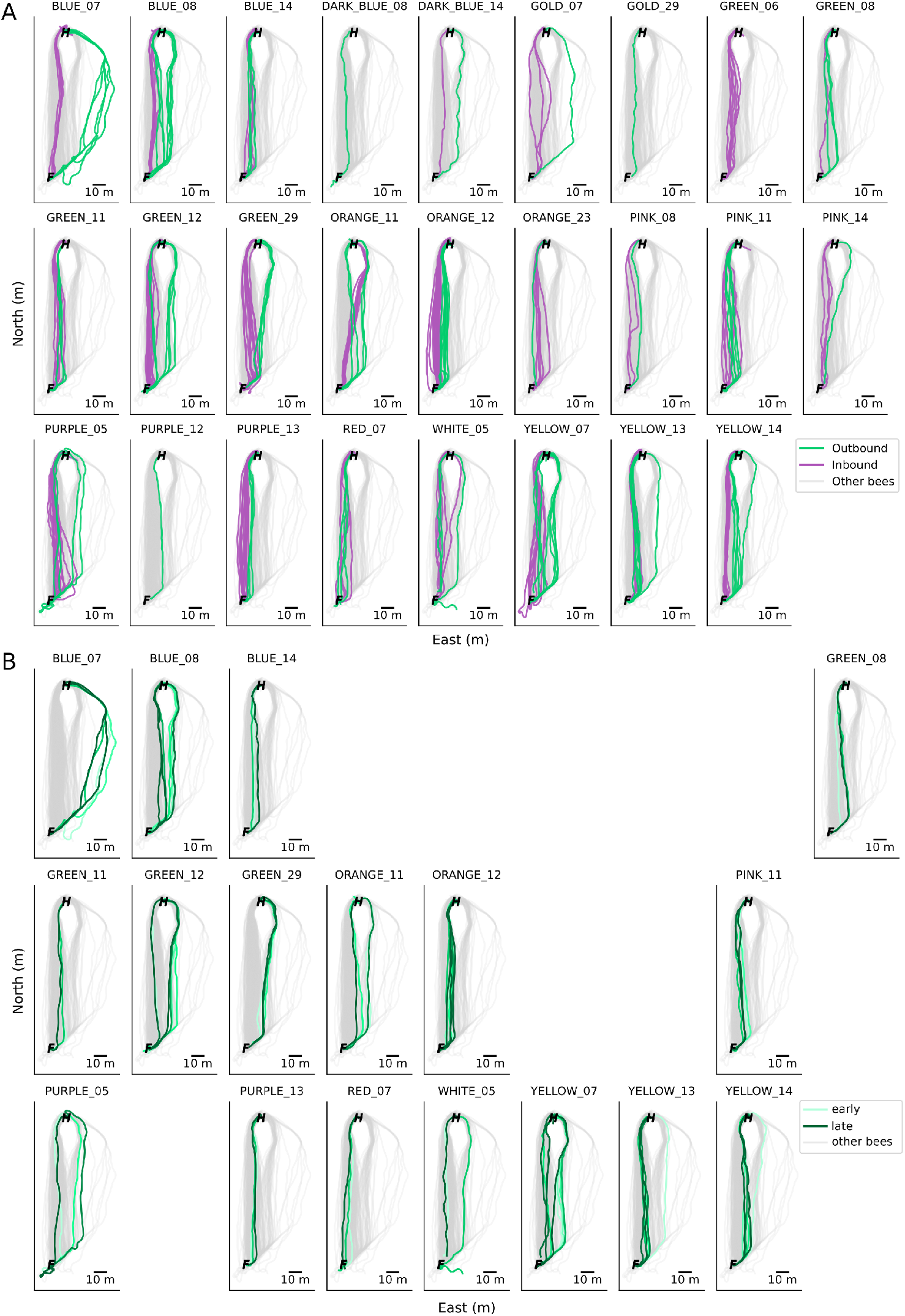
All bee trajectories between the hive and feeder included in our analyses. (a) All outbound (green) and inbound (purple) flights which reached inclusion criteria for each bee. Outbound and inbound trajectories of all other bees are also shown (grey). (b) All outbound trajectories for each bee. Trajectory order is indicated from light green to dark green. Outbound trajectories of all other bees are also shown (grey). 21

**Video 1** – 3D trajectory reconstructions for all outbound flights. Axes are as in Figure 1D and are each 10 meters in the North, East, and Up directions. Each color is one individual bee.

**Video 2** – 3D trajectory reconstructions for all inbound flights. Axes are as in Figure 1D and are each 10 meters in the North, East, and Up directions. Each color is one individual bee.

**Video 3** – Video showing pilot’s view during one flight of bee BLUE_14 from the feeder to the hive. Overlayed are the IR camera footage showing the reflector used for fast lock tracking (upper left) and high magnification camera footage showing the bee in color (upper right).

